# Mice lacking paternal expression of imprinted *Grb10* are risk-takers

**DOI:** 10.1101/2020.02.25.962399

**Authors:** Claire L. Dent, Kira D. A. Rienecker, Andrew Ward, Jon F Wilkins, Trevor Humby, Anthony R. Isles

**Author notes:** Author for correspondence: Anthony R. Isles.

## Abstract

The imprinted genes *Grb10* and *Nesp* influence impulsive behavior on a delay discounting task in an opposite manner. A recently developed theory suggests that this pattern of behavior may be representative of predicted effects of imprinted genes on tolerance to risk. Here we examine whether mice lacking paternal expression of *Grb10* show abnormal behavior across a number of measures indicative of risk-taking. Although *Grb10*^*+/p*^ mice show no difference from wild type littermates in their willingness to explore a novel environment, their behavior on an explicit test of risk-taking, namely the Predator Odour Risk-Taking task, is indicative of an increased willingness to take risks. Follow-up tests suggest that this risk-taking is not simply due to a general decrease in fear, or a general increase in motivation for a food reward, but reflects a change in the trade-off between cost and reward. These data, coupled with previous work on the impulsive behaviour of *Grb10*^*+/p*^ mice in the delayed reinforcement task, and taken together with our work on mice lacking maternal *Nesp*, suggest that maternally and paternally expressed imprinted genes oppositely influence risk-taking behaviour as predicted.

## INTRODUCTION

The imprinted genes *Grb10* and *Nesp* affect impulsive choice behavior in opposing direction^1,2^. Mice lacking paternal *Grb10* (*Grb10*^*+/p*^) prefer a larger, but delayed reward to a smaller, but more immediate reward in the delayed reinforcement task (DRT)^1^. In contrast, mice lacking maternal *Nesp* (*Nesp*^m/+^) choose a more immediate, smaller reward over a larger, but delayed reward in the DRT^2^. These behavioral findings, coupled with co-localisation of expression of *Nesp* and *Grb10* in a number of brain regions^3,4^ and cell types^1^, has led to the suggestion that they may have an antagonistic effect on the control of behavior^1,5^. This fits with the general idea that genomic imprinting evolved as a consequence intragenomic conflict between maternal- and paternally-derived alleles arising as a consequence of kin-selection^6,7^.

Recently a theoretical basis for how imprinted genes may influence risk-taking behavior has been proposed^8^. The theory comes from an extension of a model of bet-hedging, where an allele that leads to reduced mean reproductive success can be favored by selection if the allele also leads to a sufficiently large reduction in reproductive variance. An intragenomic conflict arises because the trade-off between selection on mean and variance is different for maternally and paternally inherited alleles. When reproductive variance is higher in males (as it is for most mammals) selection favors reduction of reproductive variance more strongly for paternally inherited alleles. When an allele is maternally inherited, selection more strongly favors increased mean reproductive success, even at the cost of increased reproductive variance. Following the “loudest voice prevails” principle^9^, this predicts that paternally expressed imprinted genes will promote risk-averse, variance-reducing behaviours, while maternally expressed imprinted genes will promote risk-tolerant, variance-increasing behaviours.

Wilkins and Bhattacharya suggest that the opposing pattern of behavior shown by *Grb10*^+/p^ and *Nesp*^m/+^ mice in the DRT supports this idea. The DRT is generally considered a measure of impulsive choice^10,11^; *Grb10*^+/p^ mice are more likely to choose the larger reward and are interpreted as less impulsive. However, in a naturalistic environment, delay introduces the possibility a reward will be lost to a competitor (loss of opportunity), or that the individual will be exposed to risk of predation before receipt of the reward (cost in death or injury)^12-15^. Choosing a delayed, larger reward in the DRT may therefore indicate not only less impulsive, but also more risky behavior^16^.

The idea that the choice of the more immediate, but smaller reward, displayed by *Nesp*^m/+^ mice in the DRT may reflect less risk-taking is supported by their behavior in other tasks. *Nesp*^m/+^ mice show altered reactivity to, and are less willing to explore, novel environments^3^. The propensity to explore a novel environment is regarded as a good index of risk-taking behavior and has been used as such in a number of studies^17^. However, the data for the behavior of *Grb10*^*+/p*^ mice being indicative of an increased willingness to take risk are more equivocal. Here we address this, by examining the behavior of *Grb10*^*+/p*^ mice in a novel environment. We also directly tested risk-taking behavior using the Predator Odor Risk-Taking (PORT) task. The PORT task was developed by us^18^, and has been used by others in both mice^19,20^ and rats^21^, to specifically assess “real-life” risky situations where there is a trade-off between a cost (risk of predation) and a food reward. Although *Grb10*^*+/p*^ mice show no difference from wild type animals in their willingness to explore a novel environment, their behavior on the PORT task is indicative of increased risk-taking. Follow-up tests suggest that this risk-taking is not simply due to a general decrease in fear, or a general increase in motivation for a food reward, but reflects a change in the trade-off between cost and reward.

## MATERIALS AND METHODS

### Animals

All procedures were conducted in accordance with the UK Animals (Scientific Procedures) Act 1986 under the remit of Home office license number 30/3375 with ethical approval at Cardiff University. The *Grb10* null line was maintained on an F1-hybrid B6CBA F1/crl line from Charles River. Due to potentially confounding metabolic phenotypes associated with *Grb10*^m/+^ mice ^22^, comparisons of *Grb10*^+/p^ were made with wild-type (WT) littermate controls. Subjects were male mice aged between 3-6 months during testing. All mice were housed in single-sex, environmentally enriched cages (cardboard tubes, shred-mats, chew sticks) of 2-5 adult mice per cage. Cages were kept in a temperature- and humidity-controlled animal holding room (21 ± 2°C and 50 ± 10% respectively) on a 12-hour light-dark cycle (lights on at 7:00 hours, lights off at 19:00 hours). Standard laboratory chow was available *ad libitum*, but during the progressive ratio experiment water was restricted to 2h access per day. This regime maintained the subjects at ≈90% of free-feeding body weight and motivated the animals to work for the food reward used in the task. All testing occurred during the light phase. Two separate cohorts were used. Cohort 1 (WT, n=11; *Grb10*^+/p^, n=13) undertook locomotor activity, novelty place preference, milk consumption test, PORT task, acoustic startle with a minimum of three days between each test. Cohort 2 (WT, n=14; *Grb10*^+/p^, n=11) undertook milk consumption testing and the progressive ratio, again with a minimum of three days between each test. All animals within a cohort were used throughout testing but a small number of animals were removed or lost (due to an inability to perform training stages or death) as testing progressed.

### Locomotor Activity

Locomotor activity (LMA) was measured using a battery of 12 activity cages, each measuring 21 × 21 x 36cm. The activity cages were clear Perspex boxes containing two transverse infrared beams 10mm from the floor, spaced equally along the length of the box, linked to an Acorn computer using ARACHNID software (Cambridge Cognition Ltd., Cambridge, UK). Activity was measured for two-hour sessions in the dark, over 3 consecutive days; data were collected in 5-minute bins throughout each session. Testing took place at the same time every day. The cages were thoroughly cleaned after each animal, using 1% acetic acid solution. The main measure was ‘runs’, recorded when the animal broke the two infrared beams consecutively.

### Novelty Place Preference

Novelty place preference (NPP) was assessed using an apparatus consisting of two adjacent boxes with an opening in the middle, which could be occluded by a door. These two arenas were made distinct by the colour (black or white) and the texture of the floor (smooth plastic or sandpaper). In the first stage of the test the door was closed and the animal was placed in one side of the box for one hour and allowed to habituate to that side. Then the mouse was taken out, the door removed and the mouse put back in the same side and allowed to explore both the habituated side and the novel side for a total of 30 minutes. The side to which the animal was habituated, and in which the sand-paper was placed, was pseudo-randomly allocated to avoid confounding the results. The arena was thoroughly cleaned after each animal, using 1% acetic acid solution.

The movement of each subject was tracked using a camera mounted approximately two metres above the test arena, connected to EthoVision Observer software (Noldus Information Technology, Netherlands). Behavioral measures, obtained automatically, included the duration of time spent in each zone, frequency of entries, and latency of first entrance into the novel zone. The time spent in and the number of entries into the novel compartment was measured automatically by a video tracking system, using Noldus software.

### Predator Odor Risk-Taking (PORT) task

The PORT task was conducted using the same methods and apparatus as previously described^18^ and full details can be found in the Supplementary Information. Briefly, following habituation to the apparatus, animals were trained to leave the start chamber, traverse the middle chamber, and to collect a food reward in the third chamber of the apparatus. For training trials, clean standard mouse bedding (wood shavings) was distributed evenly over the floor of the middle chamber. Trial length was set to 10 minutes but was terminated when the subject had traversed the apparatus and was observed to have collected the reward. The mouse was then removed and placed in a holding box until the start of the next trial. In the test trials the middle chamber bedding was mixed with either ‘self-odor bedding’ (bedding taken from the mouse’s home cage), or ‘predator-odor bedding’ (wood shavings mixed with a synthetic predator cue, 2,4,5-trimethylthiazoline, TMT; Contech Inc., Canada). The main measurements taken in each trial was the latency to leave the start chamber.

### Predator odor enhanced acoustic startle response (POE-ASR)

The POE-ASR was assessed in two separate test sessions, a week apart, immediately following a 10-min exposure to either untainted wood-shaving bedding (control condition) or fox odor-tainted bedding, mixed at the same concentration of TMT as used in the PORT task (see above). The order of odor presentation was counter-balanced between mice. ASR was monitored using SR-Lab apparatus (San Diego Instruments, USA), according to the previous method used^18^ (see Supplementary Information for full details).

### Condensed Milk Test

In order to increase motivation and performance of the mice in the progressive ratio task mice were placed on a schedule of water restriction. Water was maintained at a 2hr regime for the duration of the experiment, and food was available *ad libitum* at all times (apart from when in chambers). Body weight was monitored prior to water restriction, and throughout the first 10 days of restriction. Once body weight had stabilised (>90% free-feeding weight) the mice were habituated to the condensed milk reward used in the operant tasks. This was carried out using the Condensed Milk Test (CMT) to check for preference of condensed milk over water as described previously^18^ (see Supplementary Information for full details). The amount drunk and preference for condensed milk over water was measured on five successive days. This was to prevent mice having a neophobic reaction to the reward during the experiment and also to test for any differences in consumption and/or acquisition of a preference.

### Progressive ratio

All the sessions of the Progressive Ratio (PR) task were performed in 9-hole operant chambers (Cambridge Cognition Ltd, U.K) modified for use in mice, as described previously^23^. For the PR task, only the central nose-poke hole was used. The mice were presented with a visual stimulus (light) recessed into the holes and were trained to respond to this stimulus with a nose-poke as recorded by infra-red beams spanning the hole. Reward was presented in a recessed compartment on the wall opposite to the nose-poke/stimulus array. The control of the stimuli and recording of the responses were managed by an Acorn Archimedes computer with additional interfacing by ARACHNID (Cambridge Cognition Ltd). For all operant testing, animals were maintained on a restricted water access schedule, water provided for two hours immediately after testing.

During training and testing, a nose-poke in the illuminated central hole resulted in the presentation of 20 µl of an 10% condensed milk (Nestle) reward. Collection of this reward initiated a subsequent trial. Conditioned reinforcement (CRf - one nose poke required for reward delivery) was carried out for five days. Following this, a progressive ratio (PR) schedule was carried out. Here, the number of nose pokes required to receive a reward ascended linearly every four trials for three (FR4) sessions. FR4 sessions were followed by three FR2 sessions, where the number of nose pokes required to receive a reward ascended linearly every two trials. These PR sessions were followed by three CRf sessions.

A number of measures were taken, including number of rewards received, latency to first nose-poke and latency to collect the reward. During the PR sessions, an additional measure, the maximum number of nose pokes an animal was willing to make to receive a reward, was deemed the “breakpoint” and was the main indication of the animal’s motivation to work for a reward.

### Statistical analyses

All behavioral data were analysed using SPSS 20 (SPSS, USA). Data were assessed for normality and then analysed by Student’s t-test or mixed ANOVA, with between-subjects factors of GENOTYPE (*Grb10*^+/p^ vs. WT), and within-subject factors BIN; DAY; CHAMBER (start, middle or reward chamber of PORT task); ODOR (control or fox odors in PORT and POE-ASR); DAY (day of testing on CMT); SESSION (CRf, FR4 or FR2 session in PR task). For repeated-measures analyses, Mauchly’s test of sphericity of the covariance matrix was applied; significant violations from the assumption of sphericity were subject to the Huynh– Feldt correction to allow more conservative comparisons by adjusting the degrees of freedom. Preference for exploring the novel environment in the NPP test was tested using Kolmogolov Smirnov test. All significance tests were performed at an alpha level of 0.05.

## RESULTS

### Grb10^+/p^ mice show normal reactivity to novel environments

We used two measures to assess the reactivity of *Grb10*^+/p^ and WT mice to novel environments. Firstly, locomotor activity was measured in activity chambers over three consecutive days. On day 1 of testing, *Grb10*^+/p^ and WT mice showed robust levels of activity that reduced over the course of the two hour session (Figure 1A; main effect of BIN, F_23,506_=11.55, P<0.001, partial η^2^=0.344). This habituation to the environment over time was also seen over consecutive daily sessions, with the total activity levels being highest on Day 1 and reducing with consecutive daily sessions (Figure 1B; main effect of DAY, F_1.30,25.92_=7.70, P=0.006, partial η^2^=0.278). However, there were no significant differences between *Grb10*^+/p^ and WT mice either in total levels of activity across the first two hour session (Figure 1A; main effect of GENOTYPE, F_1,22_=0.48, P=0.496, partial η^2^=0.021), or the three days (Figure 1B; main effect of GENOTYPE, F_1,20_=0.004, P=0.952, partial η^2^=0.0002), or in the degree of habituation, as inferred from the rate of change in activity and indicated by the lack of a significant interaction between GENOTYPE and BIN on Day 1 (Figure 1A; F_23,506_=1.08, P=0.363, partial η^2^=0.047), and GENOTYPE and DAY across all locomotor activity sessions (Figure 1B; F_1.30,25.92_=0.42, P=0.572, partial η^2^=0.021).

**Figure 1.**
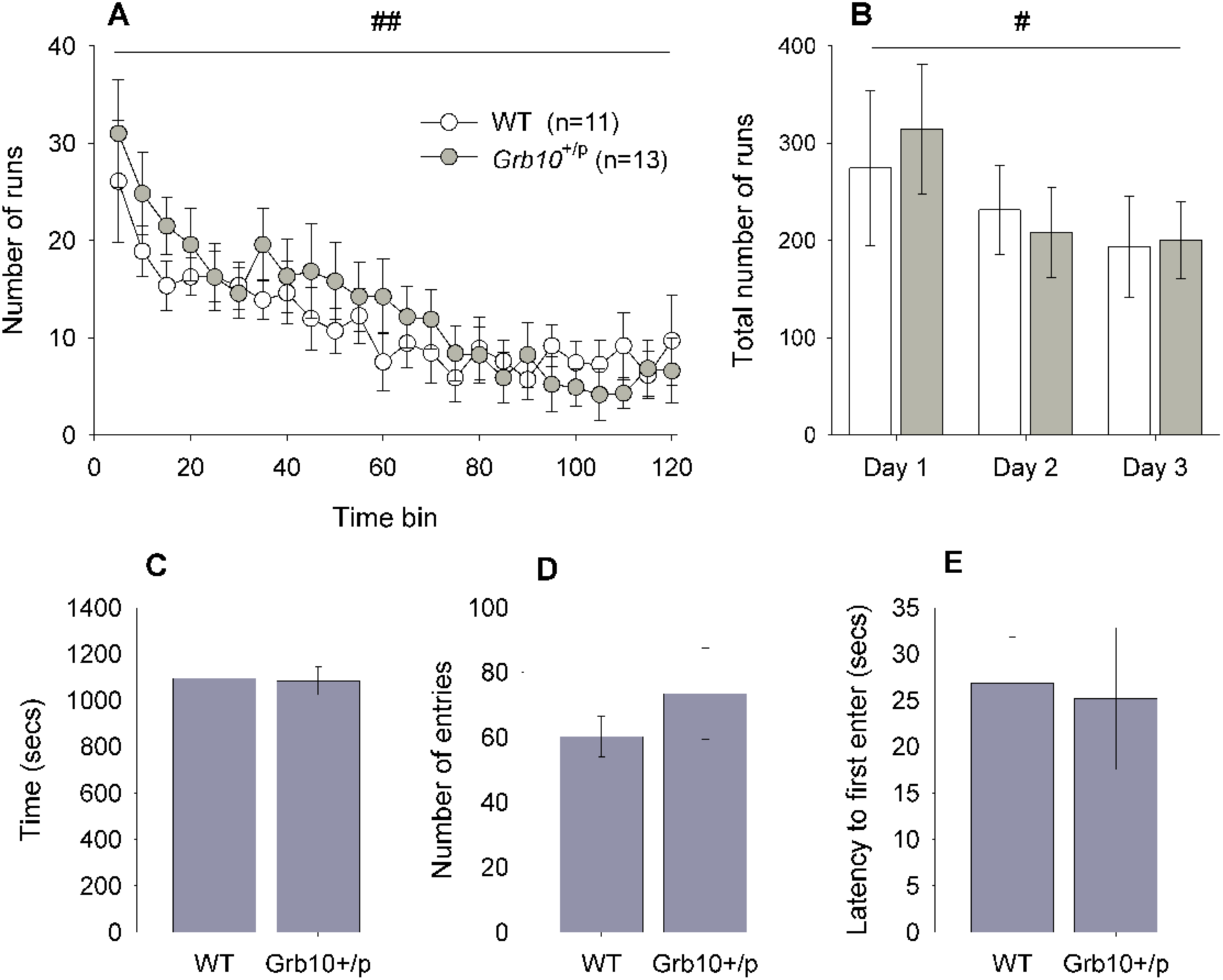
Locomotor activity and novelty place preference behavior in in *Grb10*^+/p^ and WT littermates. As the LMA session progress, activity reduces (**A**), a pattern also seen across consecutive days (**B**). However, there were no activity differences detected between *Grb10*^+/p^ and WT mice. Similarly, in the NPP test all animals showed a preference in the proportion of time spent in the novel environment, but there were no differences absolute time spent in the novel chamber between *Grb10*^+/p^ and WT mice (**C**). This was supported by other measures in the NPP test, including number of entries into (**D**) and latency to first enter (**E**) the novel environment. Data are mean values ±SEM. # (P<0.05) and ## (P<0.01) indicate within subject (factors BIN or DAY) differences.

We then explicitly measured the willingness of *Grb10*^+/p^ and WT mice to explore a novel environment using a novelty place preference test (NPP). During the test phase, both WT and *Grb10*^+/p^ mice spent significantly more time than by chance in the novel chamber (approximately 60%; Kolmongolov Smirnov test, WT P=0.003, *Grb10*^m/-^ P=0.048). Analysis of absolute measures suggested there was no distinction between *Grb10*^+/p^ and WT mice in the total exploration of the novel chamber, with no significant differences in total time (Figure 1C; t=-0.14, P=0.89), number of entries (Figure 1D; t=0.77, P=0.45) or latency to first enter (Figure 1E; t=-0.16, P=0.87) the novel chamber.

### Grb10^+/p^ mice show increased risk-taking in the PORT task

There were no differences between *Grb10*^+/p^ and WT mice during the 20-min session of habituation to the PORT apparatus (Supplementary Information, Table S1). *Grb10*^+/p^ and WT mice spent equivalent amounts of time (main effect of GENOTYPE, *F*_1,22_ = 0.85, *P*=0.37 partial η^2^=0.037) and made the same number of entries to each chamber of the PORT apparatus (main effect of GENOTPYE, *F*_1,25_ = 1.13,*P*=0.30 partial η^2^=0.049). More entries were made in the middle chamber (main effect of CHAMBER, F_1.52, 33.47_=120.2, *P* = 0.0001, partial η^2^=0.85), as might be expected as the mice traversed the apparatus, and this was also reflected in an increase in the amount of time spent in this chamber (main effect of CHAMBER, F_1.24, 33.47_=856.0, *P* = 0.0001, partial η^2^=0.98). During task acquisition, the mice were trained to cross the apparatus to collect a reward, passing through the middle chamber which had plain wood shavings on the floor (Supplementary Results Figure S1). Both *Grb10*^+/p^ and WT mice spontaneously demonstrated this behavior, and there was no difference between *Grb10*^+/p^ and WT mice (main effect of GENOTYPE, *F*_1,22_ = 3.78, *P*=0.072 partial η^2^=0.20). These data suggest that habituation and task acquisition were equivalent between *Grb10*^+/p^ and WT mice. As expected ^18,19,21^, the introduction of a predator odor into the middle chamber of the test arena significantly increased the overall latency to leave the start chamber, relative to the presence of a control odor (Figure 2a and b, main effect of ODOR, F_1,22_=14.73, P=0.001, partial η^2^=0.40). However, this effect was more pronounced in WT mice (significant interaction between GENOTYPE and ODOR (ANOVA, F_1,22_=6.75, P=0.016, partial η^2^=0.24). *Post-hoc* pairwise comparisons indicated that whilst latency to leave the start chamber was equivalent in control conditions (*P*=0.751), in the presence of the fox odor *Grb10*^+/p^ mice had significantly reduced latencies compared to WT mice (*P*=0.016), being on average 85s (60%) quicker (Figure 2b). Furthermore, whilst all but one of the 14 WT animals showed an increase in the time in the Start chamber following the introduction of fox odor, only half (5/10) of *Grb10*^+/p^ mice showed a similar slowing of the latency to leave the Start chamber.

**Figure 2.**
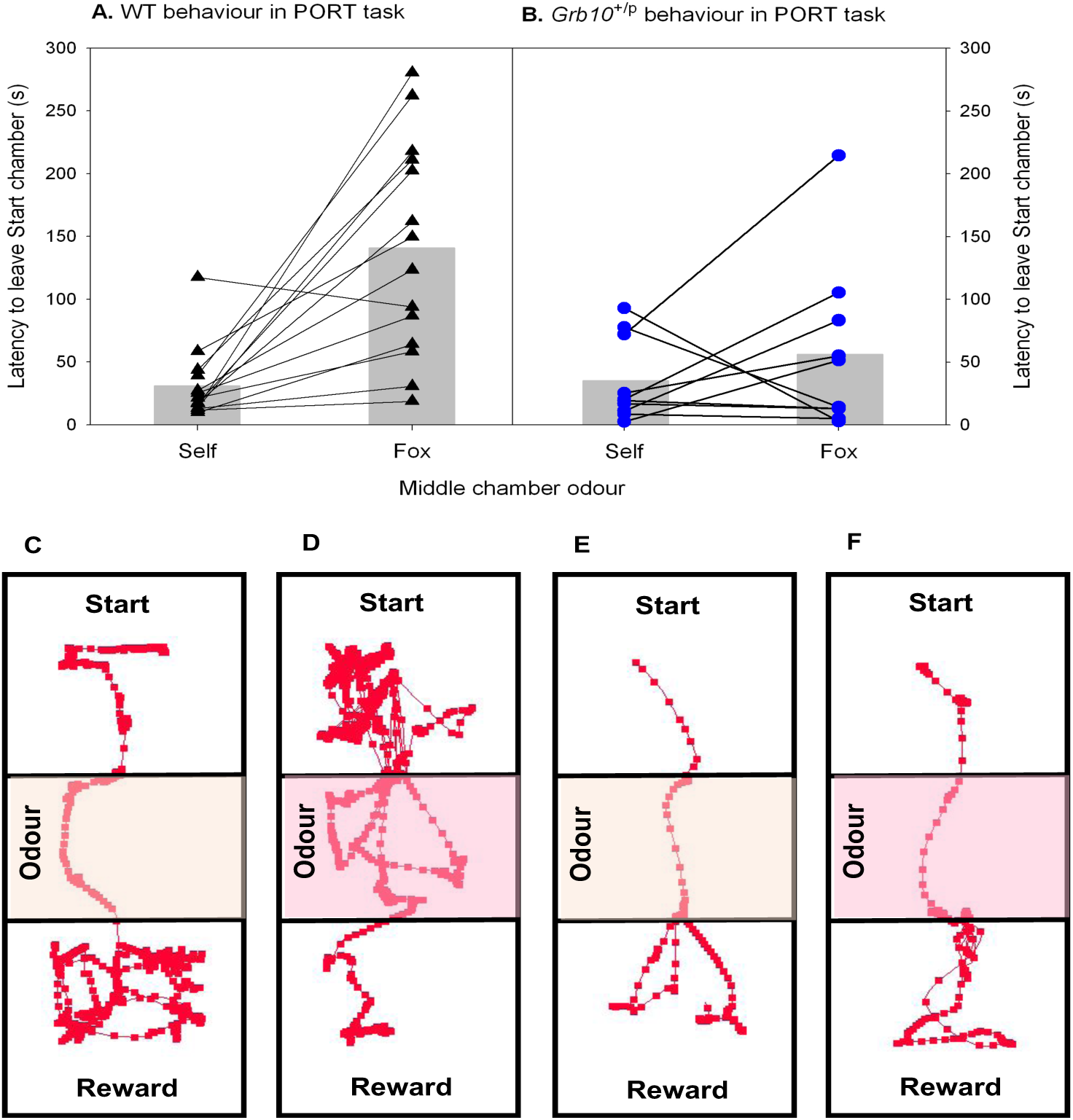
Wild type and *Grb10*^+/p^ behavior in PORT task. **A** All but one wild type animal showed an increase in latency to leave the Start chamber when a predator odor (fox) was introduced into the middle chamber of the apparatus, relative to a control odor (wood shavings from the mouse’s home cage). **B** In contrast, only 5/10 *Grb10*^+/p^ mice showed an increase upon introduction of the predator odor, and the overall magnitude of change in latency to leave the Start chamber was reduced. Representative traces from single trials of a wild type mouse with control bedding (**C**) and fox odor (**D**); and a *Grb10*^+/p^ mice with control bedding (**E**) and fox odor (**F**). Videos of these trials can be found in the Supplementary material.

As previously^18^, we confirmed that the presence of the predator odor induced an equivalent fear response in the *Grb10*^+/p^ and WT mice. Exposure to fox odor caused a significantly enhanced (24% increase) acoustic startle response (ASR) relative to prior exposure to control bedding (Figure 3a and b, main effect of ODOR, F_1,22_=11.34, P=0.003, partial η^2^=0.34), an increase that was equivalent in *Grb10*^+/p^ and WT mice (main effect of GENOTYPE, F_1,22_=2.05, P=0.166, partial η^2^=0.09).

**Figure 3.**
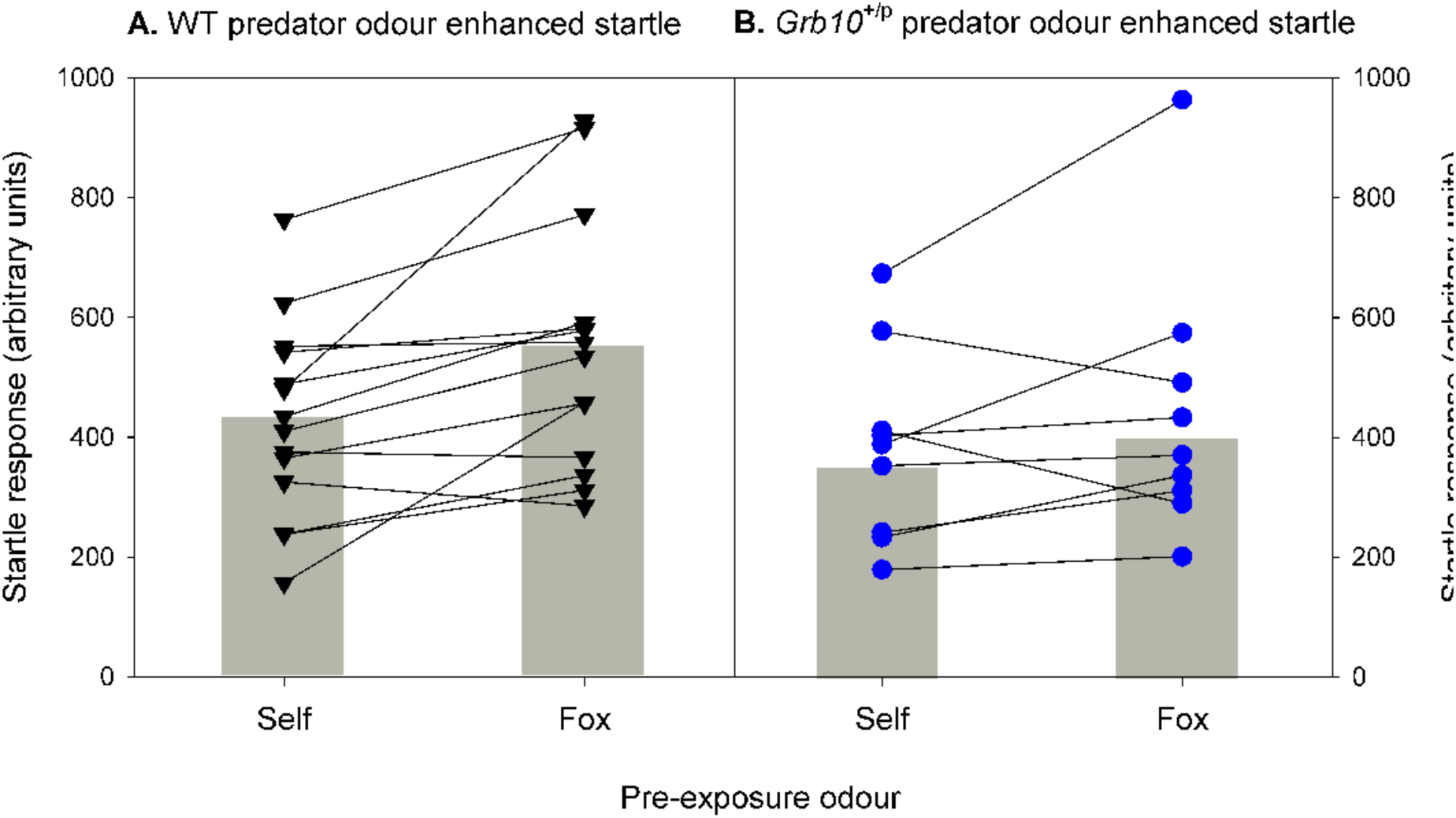
Wild type and *Grb10*^+/p^ behavior in PORT task. Acoustic startle response in both wild types (**A**) and *Grb10*^+/p^ mice (**B**) showed an equivalent increase following pre-exposure predator odor in comparison to control odor.

### Motivation for the food reward is not altered in Grb10^+/p^ mice

We performed the CMT to examine consumption and preference for a novel palatable foodstuff, namely 10% condensed milk. The total volume of milk consumed increased across successive days (Figure 4A; main effect of DAY, F_3.10,68.2_=33.28, P<0.001, partial η^2^=0.602), but did not differ between *Grb10*^+/p^ and WT mice (main effect of GENOTYPE, F_1,22_=0.34, P=0.565, partial η^2^=0.015). All animals showed an initial aversion to the novel foodstuff (preference of less than 50%), but with subsequent exposures acquired a preference for the milk reward over water (preference of approximately 75%) (Figure 4B; main effect of DAY, F_2.66,58.6_=25.63, P<0.001, partial η^2^=0.54). However, there was no difference between *Grb10*^+/p^ and WT mice in either their overall preference (main effect of GENOTYPE, F_1,22_=1.79, P=0.194, partial η^2^=0.075) nor in the rate at which their preference was acquired (interaction between GENOTYPE and DAY, F_2.66,58.6_=1.23, P=0.305, partial η^2^=0.53). These data are from the animals that went on to the progressive ratio test (Cohort 2), but a similar pattern of results was seen in animals who went on to be tested in the PORT task (Cohort 1, see Supplementary Information, Figure S2).

**Figure 4.**
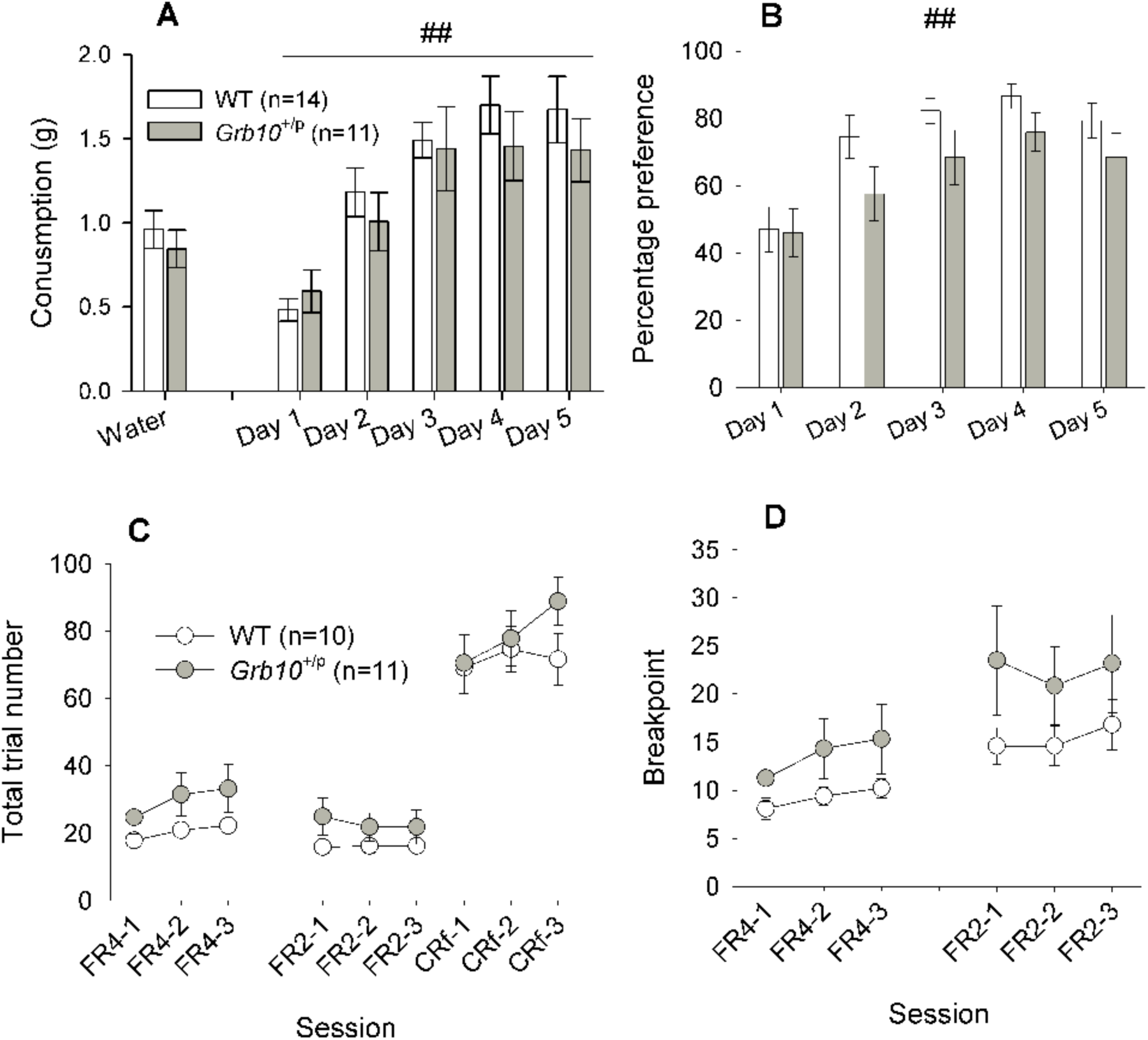
Palatable food consumption and progressive ratio behavior in *Grb10*^+/p^ and WT littermates. Consumption (**A**) and preference (**B**) for 10% condensed milk increased with successive sessions but was not different between *Grb10*^+/p^ and WT littermates. In the PR task imposition of the FR4 (number of nose pokes required to receive a reward ascends linearly every four trials) and FR2 (number of nose pokes required to receive a reward ascends linearly every two trials) reduced the total number of trials relative to CRf (one nose poke required for a reward delivery), but there was no difference between *Grb10*^+/p^ and WT littermates (C). Similarly, the BPs at FR4 and FR2 were also equivalent between *Grb10*^+/p^ and WT mice. ## indicates within subject (factor DAY) differences P<0.01.

We then examined the motivation to work for a palatable solution using a progressive ratio task. Mice were initially trained to respond on a conditioned reinforcement (CRf) schedule for 5 sessions. During the CRf sessions, subjects were able to carry out a maximum of 100 trials, which is equal to 100x 22μl rewards within each 30min test session. Analysis of the initial CRf stage revealed that both genotypes achieved the required level of performance per session, showing no effect of genotype (t_1,19_=0.80, *p*=0.44; data not shown). Following CRf training, subjects were switched to the two progressive ratio (PR) schedules: first FR4, followed by FR2. The breakpoint (BP) was defined as the maximum number of nose pokes an animal was willing to make to receive a reward and is an indication of the animal’s motivation to work for a reward. To demonstrate the effects of the imposition of the PR schedule, performance during the PR sessions were compared to the average of the three CRf sessions following PR testing (Figure 3C). Imposition of the PR schedule led to a significant reduction in the number of rewards earned within a session (main effect of SESSION, F_1.24,23.6_=160.36, P=8.80E-13, partial η^2^=0.89). There were no differences between *Grb10*^+/p^ and WT mice (main effect of GENOTYPE, F_1,19_=0.45, P=0.510, partial η^2^=0.023). This decrease in rewards earned was not due to mice running out of time to collect all the available rewards, as the average PR session did not run for the full 30 minutes and there were no significant differences in session duration between the PR (FR4 18.2mins ±1.8; FR2 19.1mins ±1.7) and CRf (21.0mins ±1.4) sessions (main effect of SESSION F_1.44,27.28_=1.36, P=0.267, partial η^2^=0.067). Although *Grb10*^+/p^ appeared to have a higher breakpoint, the main PR measure, in both FR4 and FR2 (Figure 4D), this did not reach significance (main effect of GENOTYPE, F_1,19_=1.15, P=0.296, partial η^2^=0.057). This suggests an equivalent level of motivation to work for the food reward between the *Grb10*^+/p^ and WT mice. This finding was underlined by no differences in latency measures, such as latency to first nose-poke (WT 7.97s ±1.05, *Grb10*^+/p^ 9.97s ±1.21; main effect of GENOTYPE, F_1,19_=1.56, P=0.227, partial η^2^=0.076) and latency to collect reward (WT 1.48s ±0.28, *Grb10*^+/p^ 1.84s ±0.32; main effect of GENOTYPE, F_1,19_=0.73, P=0.403, partial η^2^=0.037).

## DISCUSSION

*Grb10* is currently a unique example of an imprinted gene in which the different parental alleles show distinct patterns of expression and have distinct physiological functions^4^. We have previously demonstrated that mice lacking a paternal copy of *Grb10* have altered behavior^24^, including a higher tolerance of delayed rewards in a delay reinforcement task (DRT)^1^. One suggestion is that these changes reflect a role for *Grb10* in regulating risk-taking behavior broadly^8^. We examined this in a number of tests showing that, although *Grb10*^+/p^ mice explore a novel environment to the same extent as their wild-type littermates, they are more risk-taking on the Predator Odor Risk-taking (PORT) task. This is the direction of effects predicted from our previous analysis on the DRT^1^ and by evolutionary theory^8^, and taken together these data suggest that *Grb10* normally influences the cost vs. risk assessment and acts to reduce risk-taking.

To test whether loss of paternal *Grb10* expression would influence risk-taking we took a broad approach. Firstly, we examined the propensity of *Grb10*^+/p^ mice to explore novelty, both in terms of basic locomotor activity and habituation to a new environment, and also an explicit test of investigation of a novel environment. In both tests, the behaviour of *Grb10*^+/p^ mice was equivalent to wild-type littermates. These data were supported by the condensed milk test, which was used to assess consumption and preference for a palatable substance but can also be regarded as a measure of food neophobia. The prediction from the behavior of *Grb10*^+/p^ mice on the DRT^1^ would be that they will be more willing to take risks and therefore explore a novel environment more (or more quickly). It is possible that these tests are not sensitive enough, or that there is a ceiling effect, to detect such a ‘positive’ change. Nevertheless, it seems that across novelty domains, *Grb10*^+/p^ mice appear to behave normally.

We also used an explicit test of risk-taking to examine the behavior of *Grb10*^+/p^ mice, namely the PORT task. This task was developed by us^18^, and has been used by others to assess risk-taking in both mice^19,20^ and rats^21^. The PORT task examines a more ecologically valid aspect of risk-taking, namely the trade-off between a food reward and the risk of predation in obtaining that reward. Behavior in the task is sensitive to changes that affect this balance, such as the presence of a predator odor, or a reduction in the value of the reward^18^. In the PORT task, *Grb10*^+/p^ mice are quicker than wild-type littermates to leave the start chamber in the presence of a predator odor (fox) in order to obtain the food reward. There were no differences in habituation or acquisition of the task, and no difference in latencies in control trials, suggesting that the *Grb10*^+/p^ mice are more willing to take risks. Importantly, the difference in behavior in the PORT task is not due to changes in either fear, or motivation for palatable food alone, as predator odor enhanced acoustic startle and behavior in a progressive ratio task were equivalent between *Grb10*^+/p^ and wild-type mice. This suggests that loss of paternal *Grb10* alters the point of trade-off between a obtaining a food reward and the risk of predation in obtaining that reward.

The direction of effects in the PORT task, where *Grb10*^+/p^ mice show increased risk-taking, is consistent with findings from the DRT. Here, *Grb10*^+/p^ mice were more willing to wait for a large, but delayed, food reward. Although not developed as a direct test, behavior in discounting tasks such as the DRT have been correlated with^25^, and have been used as a proxy measure for, risk-taking^16,26,27^. Taken together, these data suggest that in the brain paternal *Grb10* normally influences the cost vs. risk decision-making, and acts to make mice more risk-averse. A complementary pattern is observed in mice lacking maternal Nesp, which show reduced exploration of a novel environment^3^ and decreased tolerance of delay in the DRT^2^. This suggests opposite effects of *Grb10* and *Nesp* on risk-taking behaviors. This suggests opposite effects of *Grb10* and *Nesp* on risk-taking behaviors. Interestingly, these genes show a strong degree of co-localisation^1^, including in neurons of the Dorsal Raphé Nucleus (DRN) and Locus Coeruleus (LC), two brain areas known to modulate risk-taking behaviours^28,29^. Moreover, these patterns are consistent with the predicted direction of effects of imprinted genes on risk-related behaviours^8^. According to this model of bet-hedging and genomic imprinting, maternally and paternally expressed imprinted genes have conflicting influences on risk-tolerance as a consequence of differences in reproductive variance between males and females. When reproductive variance is higher in males (as it is for most mammals) then paternally expressed imprinted genes like *Grb10* will promote risk-averse, variance-reducing behaviors, and maternally expressed imprinted genes like *Nesp* will promote risk-tolerant, variance-increasing behaviors.

Opposing phenotypic effects of oppositely imprinted genes is a common feature of evolutionary models of genomic imprinting. The Kinship Theory of Imprinting^30^ attributes the evolution of imprinting to an intragenomic conflict arising from differences in the inclusive-fitness effects of maternally and paternally inherited alleles. This has been most thoroughly studied in the context of pre-natal growth effects in mammals, where the fact that a female may have offspring by more than one male means that an offspring’s demands on maternal resources have a greater adverse effect on an offspring’s matrilineal kin than on its patrilineal kin. In this setting, theory predicts that paternally expressed imprinted genes will increase fetal growth, while maternally expressed imprinted genes will restrict growth, a pattern that has largely been borne out in data.

Although risk-related behaviours have not been formally modeled, the Kinship Theory has also been extended to behavioural phenotypes in a variety of ways^31-33^. It is possible that these sorts of risk-related imprinted genes could result from asymmetric inclusive-fitness effects. For example, we noted that a delayed reward can be viewed as a risky reward if there is a chance that the reward will be taken by a competitor. If that competitor is a conspecific, and the species has male-biased dispersal, the competitor would be more closely related, on average, to the focal individual’s maternal genes than their paternal genes. Those maternal genes might then favor more risk tolerance, since the fitness consequences of losing the reward would be partially offset by the indirect fitness benefit to matrilineal kin. To distinguish between these different types of explanations would require both formal modeling and a characterization of the risk-related effects of imprinted genes in a broader range of taxa. In the example outlined above, where imprinted genes affect risk due to sex-biased patterns of dispersal, we would predict risk effects to covary with dispersal patterns across taxa. The bet-hedging model^8^, which is based on sex differences in reproductive variance, would predict more consistency in the pattern of imprinted-gene effects on risk in mammals, since males have higher reproductive variance except in rare cases.

Here we test the idea that the imprinted gene *Grb10* is involved in modulating risk-taking behavior. Although not true across all domains, mice lacking paternal *Grb10* do show increased risk-taking on the PORT task. This, coupled with previous work on a delayed reinforcement task, suggest that this idea is broadly correct. Taken together with our previous work on another imprinted gene, *Nesp*, these data suggest that maternally and paternally expressed imprinted genes oppositely influence risk-taking behaviour. This “parliament of the mind”^33^, caused by opposing parental genomes pulling impulsive choice and risk-taking in different directions, is an additional factor that should be taken into consideration when considering apparently irrational or sub-optimal choice behaviors^34,35^.

## Data Availability

The data that support the findings of this study are available as Supplementary files associated with this paper.

