## Supplementary information for "Mice lacking paternal expression of imprinted *Grb10* are risk-takers"

**MATERIALS AND METHODS**

***Predator Odour Risk-Taking (PORT) task***

The PORT task was conducted using the same methods and apparatus as previously described (Dent et al., 2014) and full details can be found in the Supplementary material. For this test, the mice were placed onto a restricted home cage water schedule (2 hours access/day) for the duration of the experiment. Prior to the PORT, the mice were habituated to the reward to be used (10% condensed milk, Nestlé, UK), using previously published methods (Humby et al., 1999). In brief, mice were tested individually in cages (29 × 13 × 12 cm), during a single 10‐min session per day, across a 6‐day period (Fig. 1), with a pair of small drinking pots (3 ml volume) present at one end of the chamber. On the first day, these pots contained just water, but on the other 5 days one pot contained the reward solution (the sides pseudorandomly alternated between sessions). The volume consumed from each pot, and the reward preference was calculated for each session.

For the PORT task, the apparatus consisted of a white Perspex arena divided equally into three chambers (30 × 30 × 40 cm, depth × width × height), arranged in a row such that the middle chamber could be entered from either of the end (left/right) chambers. Guillotine doors (5 × 5 cm, width × height), operated with a pulley system, controlled access to the middle chamber. Each chamber was designated as a virtual zone for analysis (“Start”, “Middle”, “Reward”), and a small area around the reward container was also specified as an independent zone (5 cm diameter). Calculations based on the start direction (left or right) were used to determine which data would be identified as collected in the start or reward chamber, and to determine reward collection times. Reward (100μl 10% condensed milk solution) was placed in a pot attached in the centre of the reward chamber floor with Velcro.

For the PORT task, mice were initially habituated to the apparatus, and allowed to freely explore all3 chambers of the apparatus for 20 minutes (Fig. 1). For this session, the guillotine doors were open and 500 ml of clean standard mouse bedding (depth ~ 5mm) was placed in the middle chamber; there was no reward present during this session. On the next day, each subject completed five consecutive acquisition trials, in which they were placed in one of the outer chambers (‘start’), the doors were opened and the subject was free to explore the three chambers and collect the reward situated in the opposite (‘reward’) chamber. The direction of travel was kept consistent (left=start, right =reward, for example) for each day, but was counter-balanced between subjects and days of testing. For these trials, clean standard mouse bedding (wood shavings) was distributed evenly over the floor of the middle chamber. Trial length was set to 10 minutes, but was terminated when the subject had traversed the apparatus and was observed to have collected the reward. The mouse was then removed and placed in a holding box until the start of the next trial.

A similar procedure was used to assess responsivity to other odours, with three training trials in the morning with control bedding and three test trials in the afternoon with alternate odour–bedding mixtures (minimum delay between successive sets of trials was 2 hours), consisting of either ‘predator‐odour (fox odour) bedding’ or ‘self-odour bedding’ (bedding taken from the mouse’s home cage). As previously, the main parameters assessed in each trial were the latency to leave the start chamber and latency to collect the reward.

Bedding mixtures were made up fresh for each day of testing and were kept in sealed containers in a different room to prevent odour contamination. Three bedding mixtures were used: control bedding consisted of uncontaminated wood shavings; self-bedding consisted of a mixture of wood shavings with bedding material from the home cages of each subject collected on the day of testing; fox-bedding consisted of wood shavings mixed with a synthetic predator cue, 2,4,5‐trimethylthiazoline (TMT; Contech Inc., Canada), isolated from red fox (*Vulpes vulpes*) anal secretions. For the latter, a 10% TMT solution was made (in Tween), and 10 50‐μL aliquots pipetted on to 10 pieces of blotting paper (5 cm^2^), which were then placed in a bag containing 20 L of wood shavings. Therefore, every 500 ml of these wood shavings, placed in the middle chamber of the apparatus (volume 3600 cm^3^) was equivalent to 12.5 μl of 10% TMT/cm^3^, which is consistent with the concentration levels of TMT used previously to engender aversion and anxiety (Dent et al., 2014; Galliot, Laurent, Hacquemand, Pourié, & Millot, 2012).

***Predator odour enhanced acoustic startle response (POE-ASR)***

POE-ASR was monitored using a SR-Lab apparatus (San Diego Instruments, U.S.A) modified for use in mice. Animals were placed in a Perspex tube (internal diameter 35mm) and white noise stimuli were presented via a speaker. Pulse-alone trials consisted of a 40ms startle stimulus and a prepulse trial consisted of a 20ms prepulse at 4, 8, or 16db above background and a 40ms 105db startle stimulus, 70ms after the prepulse. The stimuli were presented in a pseudorandom manner every 15s. Whole body startle responses were recorded as the average startle during a 65ms window timed from the onset of the startle pulse. PPI was calculated as the percentage reduction in startle between prepulse trials and pulse alone trials.

Anxiety induced ASR [1] was assessed in two separate test sessions, a week apart, immediately following 10 min exposure to either untainted wood shaving bedding (control condition) or fox odour tainted bedding. To make the fox odour bedding 500μl 10% TMT solution (2,4,5-trimethylthiazoline (TMT), Contech Inc, Canada) was pipetted on to 10 pieces of blotting paper (5 cm^2^), which were then placed in a bag containing 20 L of wood shavings. Therefore, for every 500 ml of these wood shaving, placed into cage (volume of 3600 cm^3^) was equivalent to 12.5μl of 10% TMT/cm^3^, which is consistent with the concentration levels of TMT used previously to engender aversion, anxiety and an increased startle response [2]. The order of odour presentation was counter balanced between mice.

***Condensed Milk Test***

The CMT took place across six days, and involved putting each mouse in an individual holding box, containing two small bowls; filled with liquid. Each mouse remained in the box for 10 minutes and was free to drink from either bowl. For the first session both bowls were filled with water, to allow subjects to become familiarized with the environment, and in order to measure general water consumption. For a further five sessions, one bowl was filled with water, and the other was filled with a 10% condensed milk (Nestle, UK) solution. The positions of the solutions were alternated between left and right, to avoid location preference confounding the results. The amount of liquid drank from each container was measured after every session and the preference for condensed milk was determined. In the final session both bowls contained condensed milk, in order to ensure animals were consuming the reward.

**RESULTS**

**Table S1**

| Measure | Chamber | Wild type  Mean (±SEM) | *Grb10*^+/p^  Mean (±SEM) |
| --- | --- | --- | --- |
| Time in chamber | Start | 131 (±11.1) | 133 (9.7) |
|  | Middle | 829 (21.0) | 799 (27.8) |
|  | Finish | 100 (7.5) | 111 (10.7) |
| Entries into chamber | Start | 52 (5.2) | 61 (5.1) |
|  | Middle | 108 (7.7) | 113 (10.4) |
|  | Finish | 41 (3.6) | 52 (4.7) |

**Table S1 Measures of habituation to the PORT test data.** Measure of total time spent in, and entries to, each chamber of the PORT task apparatus during the habitation session. Both wild type and *Grb10*^+/p^ mice showed the expected pattern of behaviour [2], with no differences between genotypes.

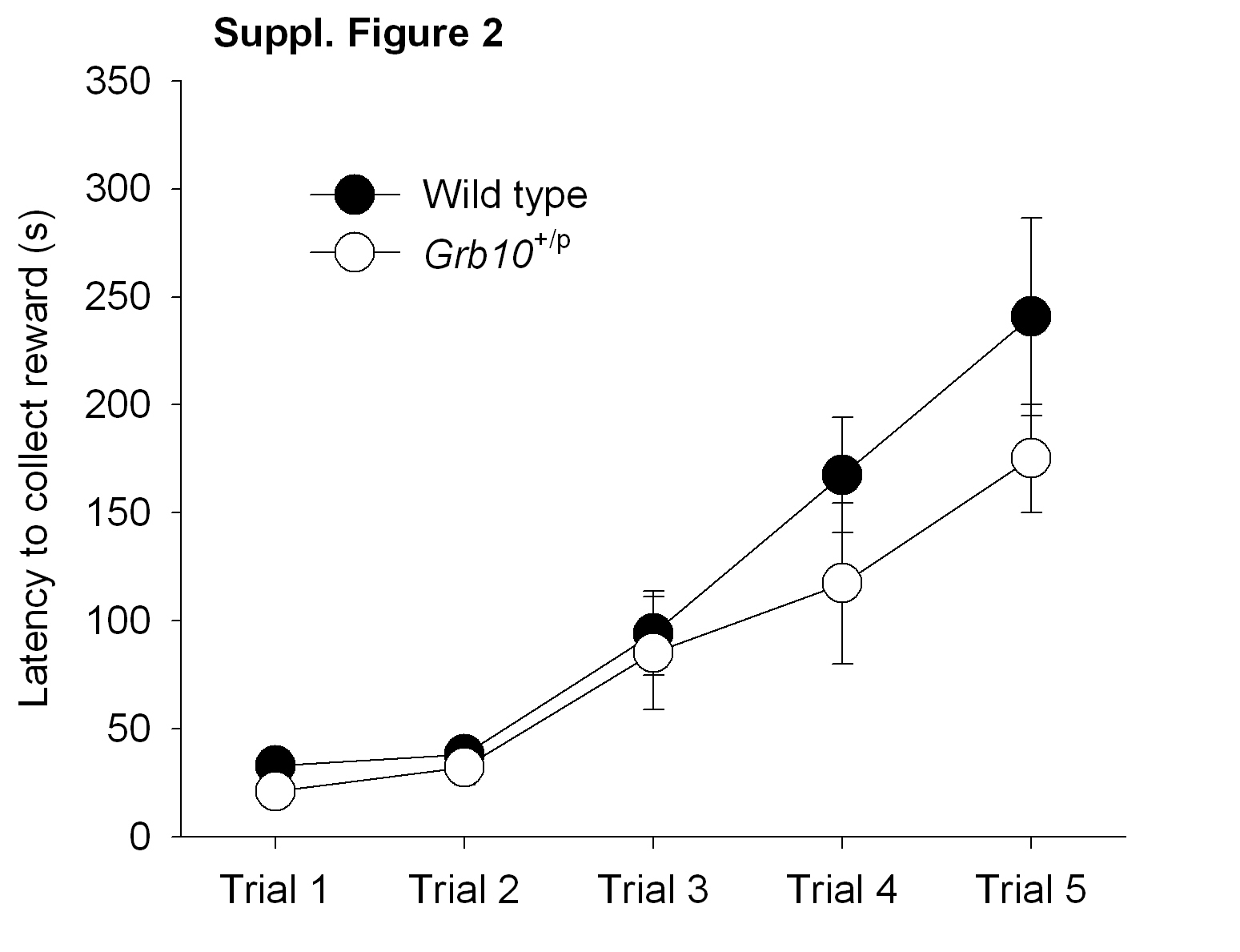

**Figure S1 Acquisition of the PORT task.** During acquisition, the mice were trained to cross the PORT task apparatus to collect the reward, passing through the middle chamber which now had clean bedding material on the floor. All the mice traversed the chambers quite rapidly. There were no significant differences between wild type and *Grb10*^+/p^ mice during these different stages of training. Graph shows means ± SEM.

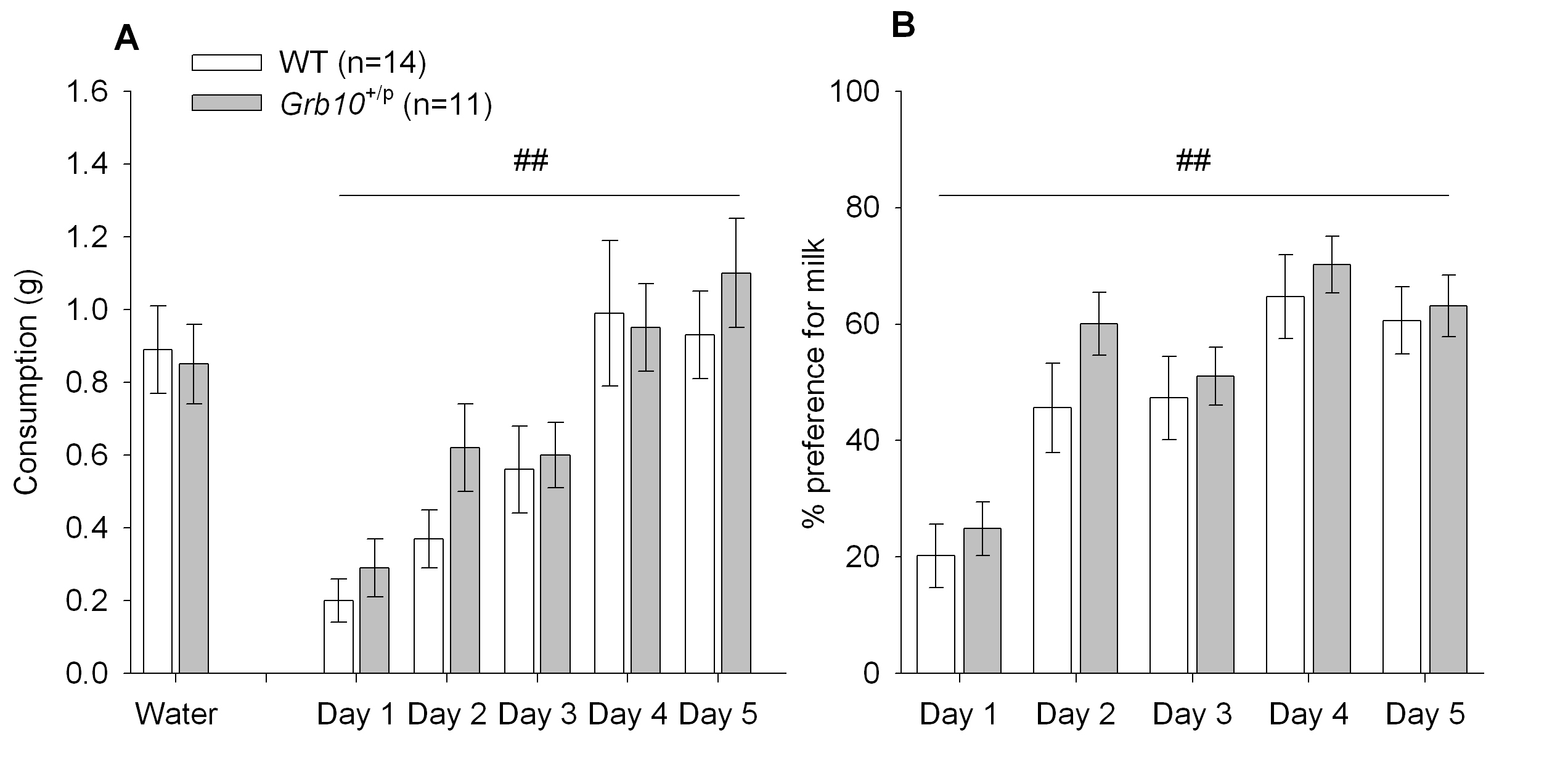

**Figure S2 Palatable food consumption in *Grb10*^+/p^ and WT littermates**. Behavior in the CMT was tested in two separate cohorts of *Grb10*^+/p^ and WT mice, these data being derived from the animals that went on to undertake the PORT task. Total volume of milk consumed (**A**) increased during the testing procedure (main effect of DAY, F_2.95,67.8_=25.5, P<0.001, partial η^2^=0.66), but did not differ between *Grb10*^+/p^ and WT mice (main effect of GENOTYPE, F_1,23_=0.59, P=0.452, partial η^2^=0.025). Similarly, preference for milk (over water) also increased with successive sessions (**B**, main effect of DAY, F_4,92_=26.7, P<0.001, partial η^2^=0.54). However, there was no difference between *Grb10*^+/p^ and WT mice in either their overall preference (main effect of GENOTYPE, F_1,23_=0.93, P=0.344, partial η^2^=0.04) nor in the rate at which their preference was acquired (interaction between GENOTYPE and DAY, F_4,92_=0.50, P=0.735, partial η^2^=0.02). Graph shows means ± SEM.
